# Transcriptomic and functional characterization indicate sexual dimorphism of discrete circadian neuron subtypes

**DOI:** 10.64898/2025.12.26.696620

**Authors:** Melina Pérez Torres, Ruihan Jiang, Nicholas Herndon, Dingbang Ma, Yerbol Z. Kurmangaliyev, Fang Guo, Michael Rosbash

**Affiliations:** Howard Hughes Medical Institute, Brandeis University, Waltham MA, 02454, USA; Department of Biology, Brandeis University, Waltham MA, 02454; NHC and CAMS Key Laboratory of Medical Neurobiology, Zhejiang University School of Medicine, Hangzhou 310058, China; Department of Neurobiology, Department of Neurology of Sir Run Run Shaw Hospital and School of Brain Science and Brain Medicine, Zhejiang University School of Medicine, Hangzhou 310058, China; Department of Molecular and Cell Biology, Harvard University, Cambridge MA, 02138; Interdisciplinary Research Center on Biology and Chemistry, Shanghai Institute of Organic Chemistry, Chinese Academy of Sciences, Shanghai, China

## Abstract

While many sexually dimorphic behaviors exhibit distinct time-of-day preferences, our understanding of how sex shapes the molecular and circuit properties of central brain neurons remain limited. Here, we uncover the transcriptomic and circuit basis of sexual dimorphism within the *Drosophila* circadian network. By leveraging single-cell RNA sequencing of male and female clock neurons, we identify specific subsets of dorsal lateral neurons (LNds), dorsal neurons 1p (DN1ps), and dorsal neurons 3 (DN3s) with dramatic dimorphic gene expression profiles. These sex differences are primarily characterized by cell-type-specific expression of genes involved in neural connectivity, particularly cell adhesion molecules (CAMs). Focusing on the dimorphic Cry-negative E3 LNds, we show that they form functionally active, synaptic connections with downstream *doublesex*-expressing pC1 and pCd-1 neurons, which serve as central regulators of dimorphic behaviors. Moreover, we demonstrate that formation and maintenance of these connections are mediated at least in part by sex-enriched CAMs, dpr9 in males and dpr3 in females. Thus, our work reveals sexual differentiation mechanisms at both the molecular and circuit levels, identifying specific molecules that sculpt sex-specific pathways to link the circadian clock to dimorphic outputs.

## Introduction

The study of circadian rhythms in *Drosophila* has focused on canonical behaviors such as eclosion and locomotor activity, revealing a core transcription-translation feedback loop (TTFL) that governs these rhythmic outputs (*1–4*). However, a wide array of additional behaviors beyond these canonical programs also exhibits pronounced time-of-day preferences, including male sex drive, female receptivity, oogenesis, and oviposition. Moreover, central brain circadian neurons, as well as components of the TTFL are directly implicated in regulating these sexually dimorphic behaviors (*5-9*).

At the heart of this regulation is the circadian circuitry, a network of approximately 240 neurons anatomically classified into seven groups: dorsal neurons (DNs-- divided into DN1a, DN1p, DN2, DN3), dorsal lateral neurons (LNds), ventral lateral neurons (LNvs) and lateral posterior neurons (LPNs) (*10*). Decades of research have identified specific roles for many of these anatomical clusters. For example, the LNvs (also known as M cells) are the master regulators of morning anticipatory locomotor activity and rhythmic activity under constant darkness (*11*, *12*). The LNds (E cells) comprise six neurons per hemisphere, subdivided into categories E1-E3, which exhibit substantial heterogeneity in both gene expression and axonal projection patterns. E1 and E2 LNds encompass four Cry-expressing neurons, which have been extensively studied for their roles in regulating evening anticipatory behavior (*11–16*). In contrast, the remaining LNds (the E3 subgroup), consisting of three Cry-negative LNd neurons per hemisphere, have remained relatively understudied due in part to their minimal connections with the rest of the circadian circuit (*14*). While DN1a and LPNs regulate daily activity/rest patterns in response to environmental temperature fluctuations, DN1p neurons are a heterogeneous population involved in sleep regulation and time-of-day perception (*17–22*). The DN2 cells regulate temperature rhythm entrainment, while DN3s primarily function as sleep-promoting neurons (*23–25*).

Recent advances in single-cell RNA sequencing (scRNA-seq) have revealed a new layer of circadian network complexity by identifying at least 27 transcriptionally unique clusters. This heterogeneity further splits the anatomically assigned cell types into multiple sub-clusters, defined by the expression of key features such as neuropeptides, GPCRs, and cell adhesion molecules (CAMs). Notably, the expression profiles of CAMs alone are sufficient to define these circadian clusters, suggesting a highly cell-type-specific expression of these connectivity molecules (*25–27*). The analysis of newly available whole-brain connectomes also revealed a matching complexity of circadian networks at the level of neuronal morphologies and synaptic connectivity patterns (*10*, *28*, *29*).

These neural connectivity molecules play essential roles in neuronal communication and wiring specificity throughout development (*27*, *30–37*). Several of these molecules have been implicated in the regulation of sexually dimorphic neuronal connectivity, such as members of the *roundabout* (*robo*) and *defective proboscis extension* (*dpr*) CAMs (*38–40*). The expression of some CAMs can be modulated by the master regulators of sexually dimorphic neuronal circuits, *fruitless* (*fru*) and *doublesex* (*dsx*). Examples include *dpr* and its connecting partner *dpr-interacting partner* (*dpr/DIP*); their expression within the fly brain identified sexual differences in cell number and neuron morphology (*40*). Multiple FruM binding sites were found in or near *dpr*/*DIP* genes, suggesting that these molecules are regulated by *fru* (*40*). FruM-specific accessible chromatin regions are more generally found near axon guidance genes and CAMs, potentially acting as sex-specific regulatory elements (*41*).

The fly brain contains multiple clusters comprising sexually dimorphic neurons. They regulate dimorphic behaviors and are characterized by sex-specific *doublesex* (*dsx*) and *fru* isoform expression (*42–45*). Among these clusters are the P1/pC1 neurons in males and pC1 in females. They are multisensory integrators of environmental cues and are necessary for successful mating. Specifically, these neurons receive chemical and auditory stimuli to execute sexually dimorphic outputs such as courtship and mate receptivity. P1/pC1 neurons interact with pCd neurons, which maintain persistent behavioral states elicited by male P1 neurons (*46*, *47*). In females, pCd neurons respond to male-specific pheromones to modulate mate receptivity (*48*). The striking time-of-day preference of several dimorphic behaviors suggests functional interactions between the circadian neurons and the dimorphic circuitry. However, the nature of these interactions and the role of sexually dimorphic gene expression within circadian neurons remain largely unexplored.

In this study, we investigated the transcriptional profile of circadian neurons with a focus on sexually dimorphic gene expression. By performing single-cell RNA sequencing of male and female circadian neurons, we identified subsets of LNds, DN3s, and DN1p neurons with sexually dimorphic differences, which are enriched for genes impacting neural connectivity, including CAMs, GPCRs, and neuropeptides. Focusing on the sexually dimorphic circadian subgroup of LNds, we show that they reside upstream of and connect to *dsx*-expressing pC1 and pCd neuronal subsets. We further demonstrate that this connection is mediated, at least in part, by sex-specific CAMs expressed within the LNds.

## Results

### Four circadian cell types show a sexually dimorphic transcriptomic profile

Our previous work leveraged single-cell RNA sequencing to characterize the transcriptomes of circadian neurons, demonstrating that they segregate into at least 27 distinct cell types (*25–27*). Given prior evidence of sexually dimorphic cell types within the circadian circuit (*49–51*), we re-examined these neurons using single-cell sequencing libraries prepared separately from male and female circadian neurons (Fig. 1A).

**Figure 1.**
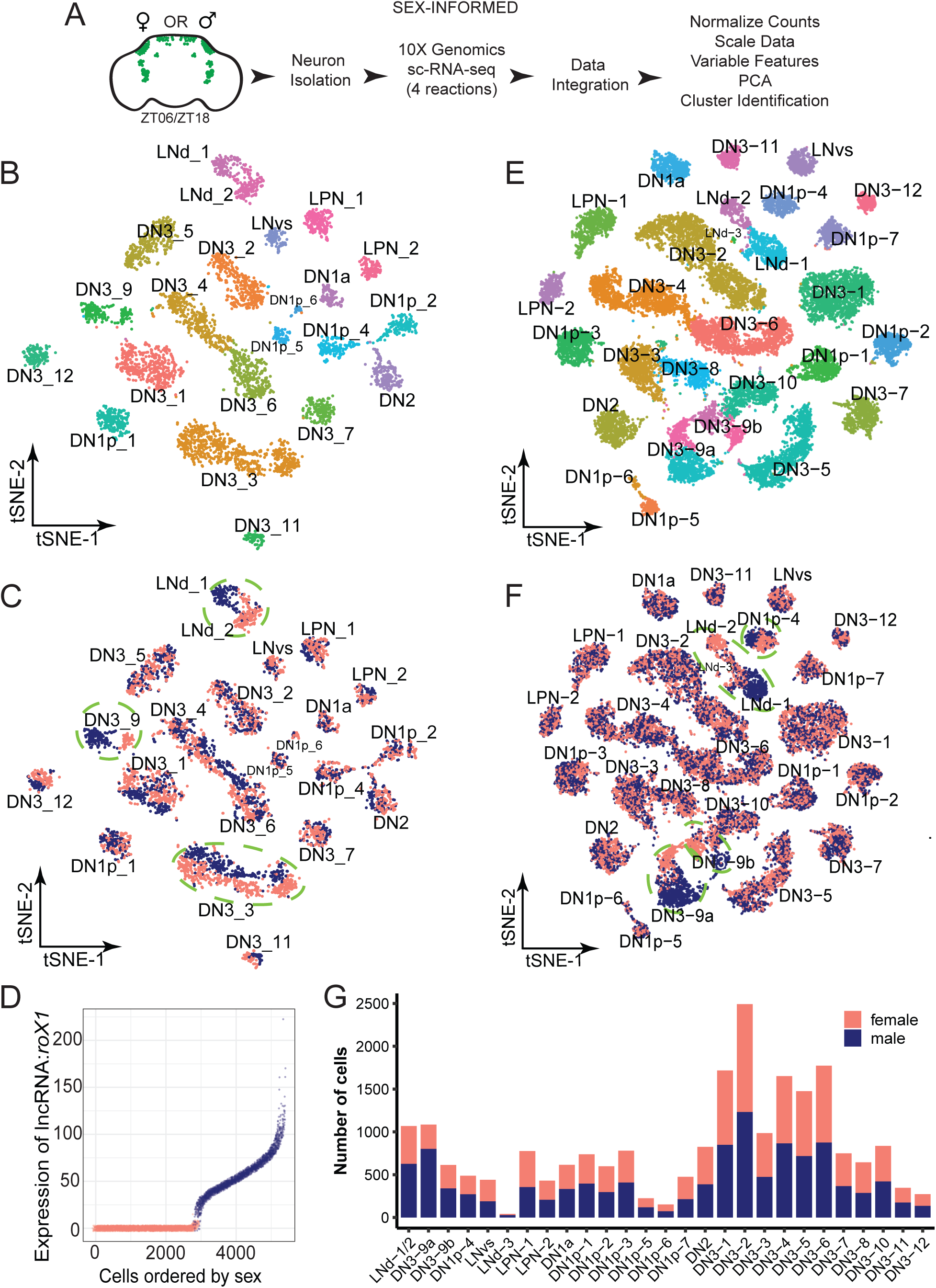
Identification of sexually dimorphic circadian transcriptomic clusters. **(A)** Schematics of experimental design for generating and processing single cells RNA sequencing of male and female circadian neurons. Male and female flies with GFP labeled circadian neurons were collected at two time points (ZT06 and ZT18). Male and female samples were collected and handled separately. GFP-positive circadian neurons were isolated using FACS, and single-cell RNA sequencing libraries were generated using single-cell 3’ RNA seq (10X Genomics). Genome alignment and data processing was carried out using CellRanger and Seurat packages, respectively. **(B)** Dimensional reduction plot of the de novo clustering identities from the newly generated sex-informed circadian single cell data set. Each color represents a distinct circadian cell type. **(C)** Dimensional reduction plot of the circadian clusters grouped by sex. Green dashed lines show sex-biased circadian transcriptomic clusters. **(D)** Normalized expression plot of *lncRNA:roX1*. X axis shows high confidence circadian neurons; pink dots show the female-derived cells and blue dots show the male derived cells. Y axis shows *lncRNA:roX1* normalized expression from the sex-informed dataset. **(E)** Dimensional reduction plot of the integrated sex-informed and sex-inferred datasets. Each color represents a distinct circadian cell type. **(F)** Dimensional reduction plot of the integrated sex-informed and sex-inferred datasets grouped by inferred sex based on ROC determined cutoffs for optimal sex-calling. Green dashed lines show sex-biased circadian transcriptomic clusters. **(G)** Number of male and female cells in integrated data set plotted by cell type. Pink bars show number of female cells; blue bars show number of male cells per cluster. Brackets indicate the sex-biased clusters.

To this end, male and female flies expressing an enhanced GFP (eGFP) fluorescent marker in all circadian neurons (*Clk856-Gal4, DN3-spGal4, UAS-eGFP*) were collected at two circadian time points (ZT06 and ZT18). Brains were dissected, and GFP-positive neurons were isolated via fluorescence-activated cell sorter (FACS). Single-cell transcriptomic profiles of male and female clock neurons were generated using 10X Genomics platform (see Methods), yielding 2,451 male and 2,865 female cells (Fig. 1B). These datasets were integrated (hereafter referred to as the “sex-informed” dataset) and used for unsupervised clustering. The results of this analysis were highly congruent with our previous studies (Fig. S1A; (*25*). Notably however, there were several cell types in which male and female transcriptomes were sufficiently different to form distinct clusters subdivided by sex (Fig. 1C, cells circled in green: blue are male, and pink are female). These data indicate that at least one cluster of LNds and two DN3 cell types exhibit pronounced sexually dimorphic gene expression patterns.

Previous studies have utilized the expression of sex-specific transcripts to divide single-cell transcriptomes into male or female cells (*50*, *52*). To determine if this approach could provide robust validation for *Drosophila* circadian neurons, we examined the expression of several sex-specific genes (Fig. 1D; Fig. S1B-D). The male-specific long non-coding RNA *roX1* was one of the most sex-specific markers and was detected in 48% of cells; male-derived cells exhibit high *roX1* transcript levels, whereas female-derived cells show virtually undetectable *roX1* expression (Fig. 1D; Fig. S1E-F). Using the Receiver Operating Characteristic (ROC) as a performance metric for true and false positive rates, we identified a high correlation (AUROC= 0.998) between *roX1* expression levels and male sex-assignment. This analysis determined that the optimal cutoff for sex assignment was 9.4 transcripts-per-10,000 UMI (TP10K) (Fig. S1G).

To increase the resolution and ensure sufficient coverage of rarer circadian cell types, we generated an additional circadian neuron transcriptomic dataset and combined it with the sex-informed dataset and a previously published circadian single-cell dataset (see Materials and Methods, Fig. 1E, Fig. S2-3) (*25*). This final data set consists of 22,306 high-confidence circadian neurons from both sexes. To infer sex in this data set, we used the sex-assignment cutoffs described above, *roX1* expression greater than 10 define male cells and less than 2.3 define female cells. In total, we assigned sex to 11,434 male cells and 10,872 female cells (hereafter referred to as the “sex-inferred” dataset, Fig. 1F-G, S2A-B). This analysis clarified a key point: two previous distinct circadian LNd clusters/cell types (9:LNd_NPF/12:LNd_AstC in Ma et al., 2021 (*26*), labeled as LNd_1/LNd_2) are in fact the same cell type separated by sex. Expression of known marker genes indicates that these sexually dimorphic clusters correspond to the heretofore understudied E3 LNds (labeled as LNd_c in the connectomes (*10*, *29*)). We also identified two closely related DN3 clusters (DN3-9a and DN3-9b) and one subset of DN1ps (DN1p-4) with dramatic sex-biased transcriptional clustering.

There are also differences in the number of male and female circadian neurons in the DN3s. The ratio of male-to-female cells within each cluster was calculated from the sex-inferred dataset, where male and female cells were processed together (Fig. S2C). Unlike the sex-informed dataset, this integrated dataset lacks cell number bias stemming from differential cell recovery. The dimorphic DN3 cluster (DN3-9a) shows three times more male cells than female cells, whereas its transcriptionally related DN3-9b cluster shows equal numbers of male and female cells. This observation is consistent with a recent study reporting more DN3s in male adult brains than in females (*51*). The sex-biased cell numbers appear more prominent for the DN3a/d clusters, as the rest of the cell types show similar if not identical numbers of male and female circadian neurons (Fig. S2C). Minor differences observed in other clusters are modest and likely reflect expected variability in cell capture and recovery, which can be most pronounced in smaller clusters.

### Sexually Dimorphic Gene Expression in Circadian Clusters is Cell-Type-Specific

To identify the molecular basis of this dimorphism, we analyzed marker gene expression within the different circadian clusters. *fru* is upregulated in these four sex-biased clusters; *fru* expression has been previously reported in a small subset of DN1s and LNds (Fig. S4A) (*50*). Although *fru* expression is detected in all circadian clusters, the dimorphic clusters express consistently higher levels of *fru* than the rest of the clusters; similar observations have been reported previously (*56*). The two sex-biased DN3 clusters also express *zelda* (*zld*), a transcription factor involved in sex determination (Fig. S4B). Moreover, a defining feature of the male LNd cluster (LNd-1) is the *neuropeptide F* (*NPF*) (Fig. S4C); previous work identified NPF as sexually dimorphic (*57*).

To further characterize sexually dimorphic features within this data set, differential gene expression analysis was performed between male and female cells from each circadian cluster. The initial comparison was conducted on the combined dataset to maximize statistical power, given that the differences between male and female cells from sex-informed datasets may be confounded by technical artefacts due to separate processing of the four samples. Therefore, we also repeated differential gene expression analysis using only the sex-inferred datasets.

The results of both these analyses were highly concordant (data not shown). Most clusters have fewer than 10 sex-differentially expressed genes, whereas four clusters manifest more significant transcriptional differences (adjusted p-value < 0.05, fold-change ≥ 1.7) (Fig. 2A). The LNd clusters show 42 male-enriched genes and 19 female-enriched genes. The two DN3 clusters (DN3-9b and DN3-9a) have 35 and 33 male-enriched genes, whereas females have 32 and 31 female-enriched genes, respectively. Lastly, the one sexually dimorphic DN1p cluster (DN1p-4) has 12 male and 17 female-enriched genes, which may be due to the smaller size of this cluster (Tables 1 and 2). The high degree of cell-type-specificity, with minimal overlap among sex-specific genes, suggests that these genes and neurons regulate distinct aspects of sexually dimorphic outputs (Fig. 2B-C).

**Figure 2.**
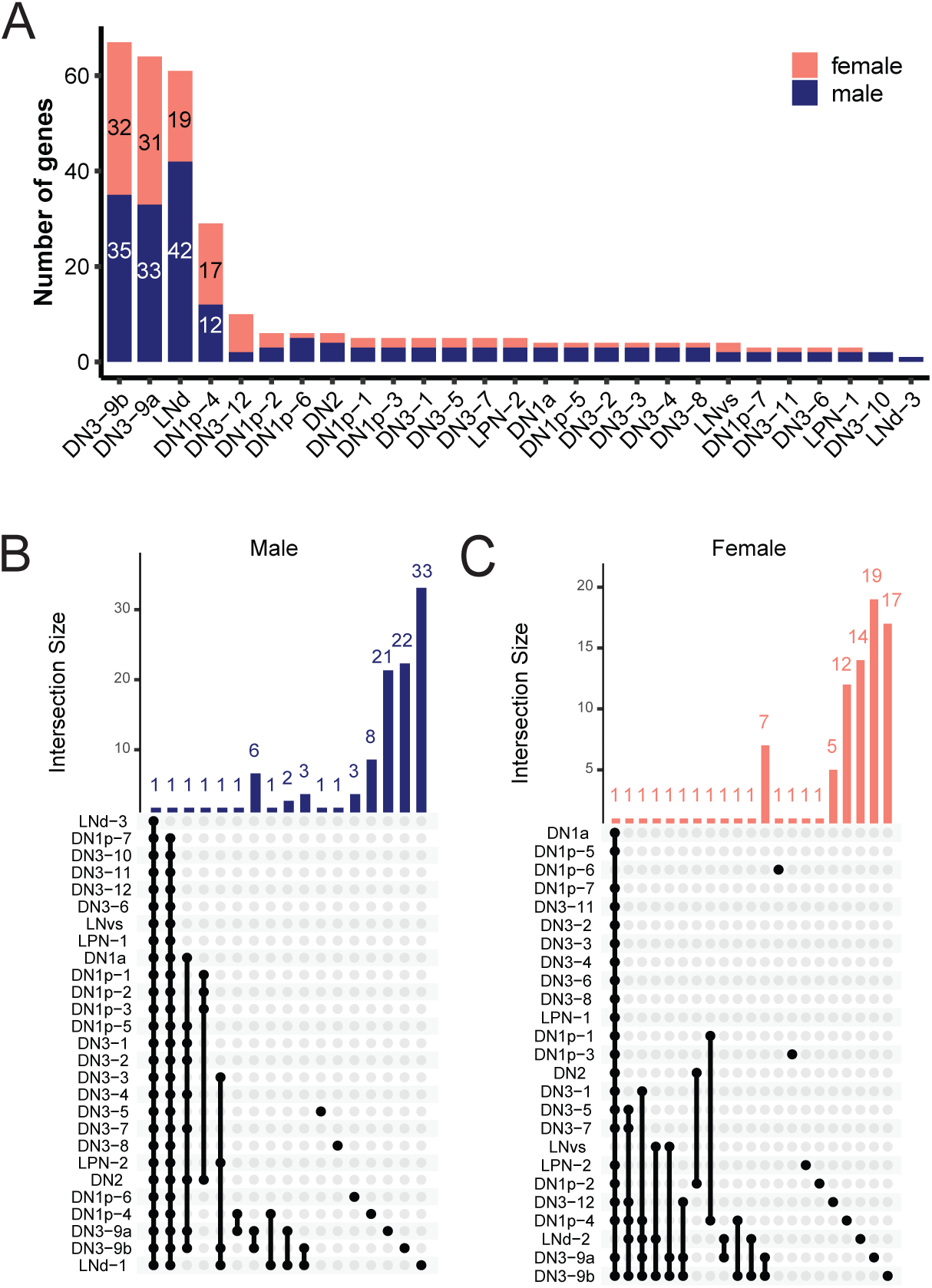
Sexually dimorphic gene expression of circadian neurons is cell-type-specific. **(A)** Bar plot showing number of differentially expressed genes (DEGs) between males and females for each individual cluster. Pink bars show number of female DEGs, blue bars show number of male DEGs. **(B)** Gene intersection plot of all genes upregulated in male circadian neurons in each cluster. Horizontal blue lines show to the gene set size, which represents the total number of male upregulated genes in each cluster. Black dots and vertical black lines show an intersection of common genes between the clusters. The vertical blue line shows the number of common genes for each identified intersection. **(C)** Gene intersection plot of all female upregulated genes in each cluster. Horizontal pink lines show the gene set size, which represents the total number of female upregulated genes in each cluster. Black dots and vertical black lines show an intersection of common genes between the clusters. Vertical pink line shows the number of common genes for each identified intersection.

As expected, *lncRNA:roX1* and *lncRNA:roX2* are exceptions and commonly upregulated in all male cells (*58*, *59*). *sister-of-sex-lethal* (*ssx*) is also significantly upregulated in several subsets of male neurons; *ssx* is known to promote male-specific features (*60*). Two additional novel sex-specific genes are encoded on the X chromosome: *Muc14a* is upregulated in nine male dorsal neurons, and *terribly reduced ocelli* (*trol*) is upregulated in male cells from three DN1p and DN2 clusters (Fig. 2B, Table 1).

The most common female-enriched gene is *Sex lethal* (*Sxl*), primary master switch during sex determination (*61*). Interestingly, the *pigment-dispersing factor receptor* (*Pdfr*) is differentially upregulated in female LNvs, LNds, and DN3-9b; notably, whole-body *Pdfr* mutants have sexually dimorphic behavioral differences (*62*). Additional differentially expressed genes in female cells include *CG15765*, identified in a genome-wide association study (GWAS) as a candidate for modulating neural input affecting sperm competition in females (*63*). Other common upregulated genes in female circadian neurons that have not yet been associated with sex-specific physiology or behavioral functions include *CG34247*, *alpha-Man-Ia*, *CG44422*, and *PVRAP* (Fig. 2C, Table 2).

### Sexually Dimorphic Circadian Clusters Are Enriched for Neural Connectivity Molecules

Due to the relatively small number of differentially expressed genes identified within individual clusters, gene ontology (GO) analysis was performed on the union of sex-differentiated genes from the top four dimorphic clusters for both sexes. This set of 101 male-enriched genes and 79 female-enriched genes were compared to a background of 3,741 genes detected in all circadian neurons. The top GO term in both sexes was the GPCR signaling pathway (GO:0007186), which includes both GPCRs and neuropeptides. Other enriched categories included GO terms related to cell signaling pathways and cell adhesion molecules (Fig. 3A-B). The same GO categories in both sexes suggest similar functions of sex-enriched genes in male and female clock neurons.

**Figure 3.**
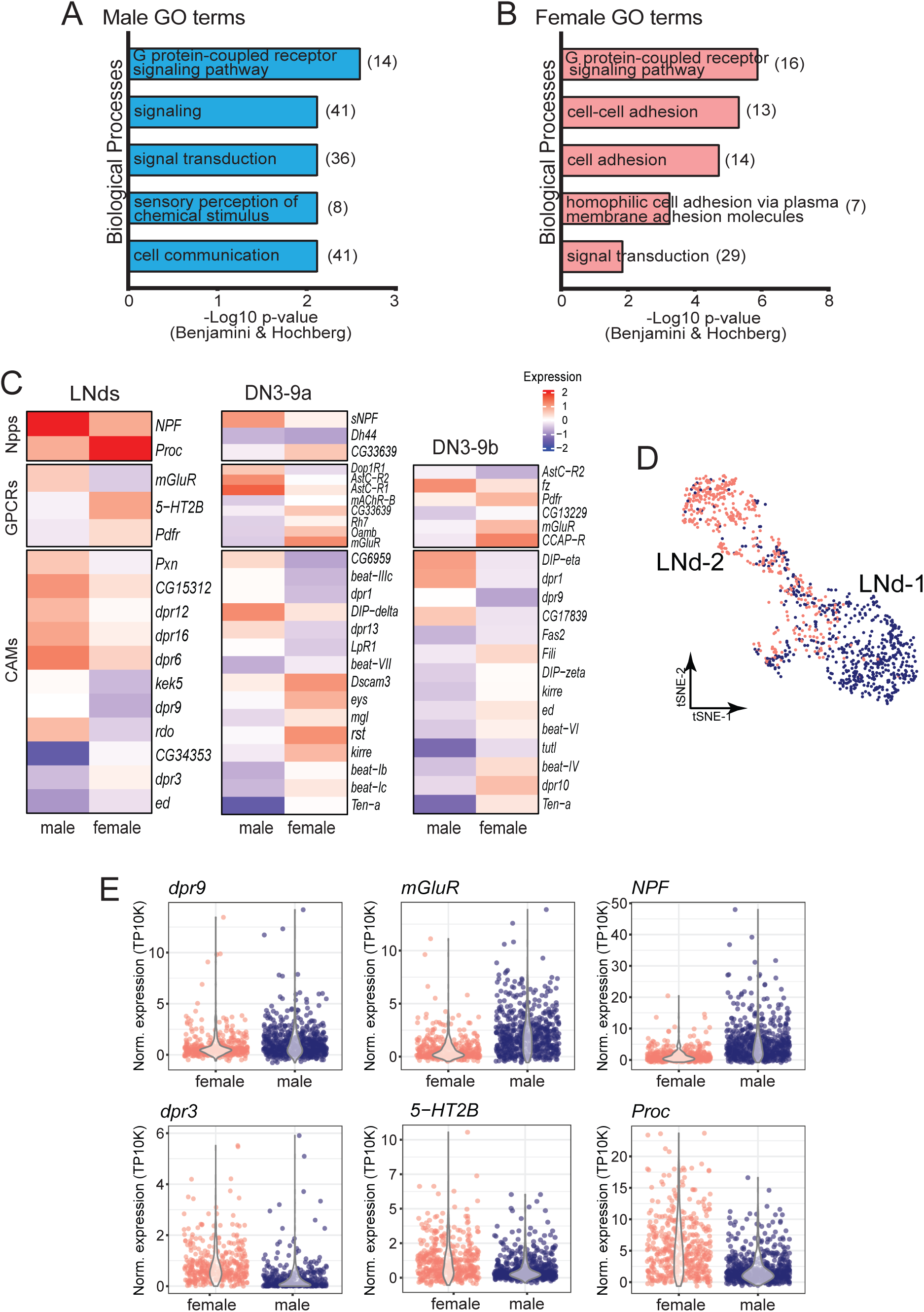
Sex-specific connectivity molecules are expressed in the sexually dimorphic circadian neurons. **(A-B)** Gene ontology analysis of **(A)** male and **(B)** female enriched genes in the sexually dimorphic circadian cell types. X axis shows –LOG10 p-value of each enriched gene group identified. Numbers in parenthesis show the number of genes identified in the dataset that form part of the gene ontology term (Benjamini & Hochberg p-value less than 0.01). **(C)** Heatmap visualizing gene expression (normalized Log2 scale) of sexually dimorphic gene groups in the LNds (left), DN3-9a (middle) and DN3-9b (right) clusters. **(D)** Dimensional reduction plot of the male (blue) and female (pink) cell distribution of the sexually dimorphic LNd cluster. **(E)** Normalized expression of neural connectivity molecules enriched in either male (top) or female (bottom) LNds. From left to right: CAMs, GPCRs and neuropeptides.

These GO categories align with the finding that neural connectivity molecules are defining features of circadian cell types (*27*): neuropeptides, GPCRs, and CAMs were indeed among the top sex-enriched genes in the four sexually dimorphic clusters. A subset of select top differential genes are shown at the single-cell level as violin plots (Fig. 3C-E, Fig. S5). The LNds show strong sexual dimorphic expression of *NPF* and *Proctolin* (*Proc*); *NPF* is enriched in male LNds, consistent with previous studies (*57*), whereas *Proc* is enriched in female LNds. Although *Proc* serves as a signaling factor important for oviposition in other cell types (*64*), this is the first report of its sexually dimorphic gene expression within circadian neurons. Three neuropeptides are dimorphically expressed in the DN3-9a cluster: *sNPF*, *Dh44,* and *CG33639*. Only *Dh44* has been reported to contribute to sex-specific behaviors in other cell types (*65*).

Although some of the sex-enriched GPCRs have been previously shown to contribute to sexually dimorphic behaviors, including activity/rest periods, oogenesis, courtship, and mating (*9*, *17*, *46*, *66–69*), the specific roles of these receptors in clock neurons is unknown. An exception is the expression of *mGluR* in subsets of LNds; it regulates the sexually dimorphic behavior of daytime activity (*17*). *mGluR* is upregulated in male sex-dimorphic LNds, and in both female sex-dimorphic DN3 clusters (DN3-9a and DN3-9b). This suggests context-dependent roles of this GPCR in dimorphic outputs based on cell identity (Fig. 3C).

The largest category of sexually differentiated genes in the dimorphic clusters are CAMs, mainly belonging to the immunoglobulin superfamily (IgSF) of proteins; they include Dpr/DIP and Beat families of cell adhesion molecules (*70–73*). Interactions between Dprs and DIPs have been shown to regulate wiring specificity of neural circuits (*74–76*). The LNd clusters express four male-enriched and one female-enriched *dpr*, with the largest differences observed for *dpr9* in male LNds and *dpr3* in female LNds (Fig. 3E). Each DN3-9 cluster expresses two male-enriched dprs; *dpr1* is shared between these two male clusters and is known to gate the timing of male courtship steps in other cell types (*38*). Overall, we have identified a significant overrepresentation of neural connectivity molecules among genes with sexually dimorphic expression patterns in specific circadian cell types.

### Sexually Dimorphic CAMs Mediate Functional Connectivity between LNds and Downstream Dimorphic Circuits

Connectome-defined LNds are subdivided into Cry-expressing LNd_a/b neurons and Cry-negative LNd_c neurons, the latter corresponding to the LNd_1 and LNd_2 transcriptional clusters in our dataset. LNds have been previously linked to various sex-specific behaviors (*13*, *77–79*), and the *DvPDF*-Gal4 driver, provides genetic access to a subset of Cry-expressing LNds (LNd_a) as well as all Cry-negative LNds (LNd_c). To address possible synaptic connectivity between LNds and *doublesex*-expressing (*dsx*) neurons known to regulate sexually dimorphic physiology and behavior (*44*, *47*, *80*, *81*), we used a modified version of the *trans-*synaptic labeling tool, *trans-*TANGO, to restrict postsynaptic labeling to neuronal subsets of interest (*82*). This restricted *trans-*TANGO uses a membrane-bound ligand expressed in axonal projections of presynaptic neurons, combined with a ubiquitously expressed QS transcription factor flanked by FRT sites for conditional expression in specific candidate cells. The *DvPDF-Gal4* driver was used to direct ligand expression in presynaptic LNd circadian neurons, and postsynaptic target neurons were restricted to *dsx*-LexA-expressing neurons via *LexAop-FLP* (Fig. 4A) (*24*).

**Figure 4.**
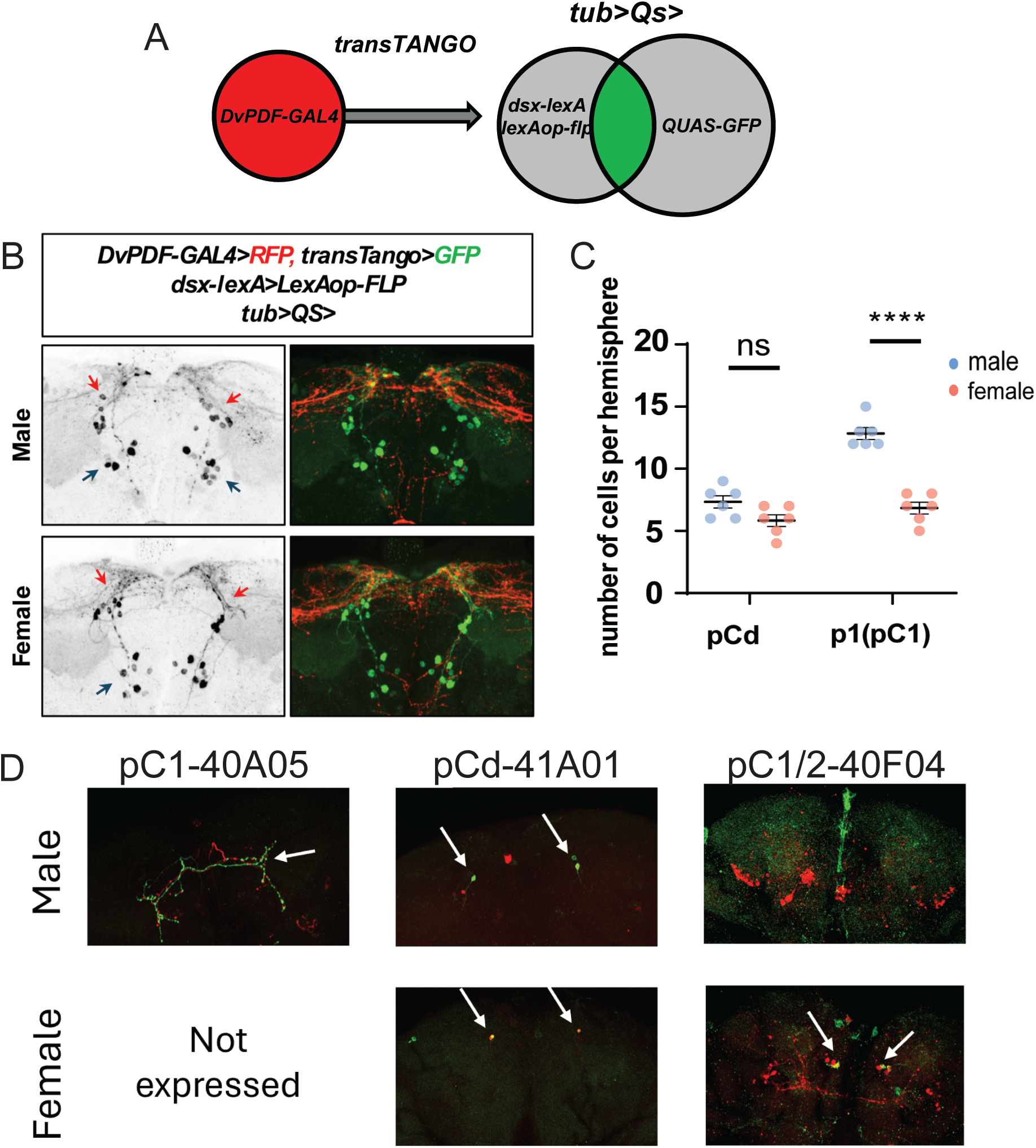
Circadian LNds are upstream of dsx-expressing pC1 and pCd neurons. **(A)** Schematic representation of restricted transTANGO approach to identify potential synaptic connections between two neuronal cell types. Gal4-labeled neurons drive the expression of transTANGO presynaptic ligand. LexA-labeled neurons drive the expression of a LexAop-FLP. This allows only the LexA-expressing neurons to flip out a STOP codon and promote the expression of the TF QS, which, in turn, promotes the expression of GFP (QUAS-GFP) in downstream neuronal subsets of interest. **(B)** Restricted transTANGO of male and female brains between *DvPDF-Gal4* and *dsx-LexA*. RFP is expressed in all *dsx-LexA*-expressing neurons, GFP is expressed in *dsx-LexA*-labeled neurons, which are post-synaptically connected to *DvPDF*-expressing neurons. Red and black arrows show the pCd and pC1 neuronal subsets, respectively. **(C)** Quantification of the number of detected dsx-expressing neurons downstream of *DvPDF*-expressing neurons. pC1 and pCd neuronal subtypes were identified by anatomical location and projection patterns. **(D)** Restricted transTANGO of male and female brains between *DvPDF-Gal4* and three sparse LexA drivers labeling (left) pC1-40A05, (middle) pCd-41A01, and (right) pC1 and pC2-40F04 dsx-expressing neuronal subsets. 40F04 male samples did not show GFP labeling. Female brains from 40F04 shows pC1 neurons labeled in green. GFP is expressed in dsx- LexA-labeled neurons which are post-synaptically connected to DvPDF-expressing neurons. White arrows show GFP-positive dsx-expressing neuronal populations.

We observed an average of 15 pCd and 25 pC1 neurons in male brains compared to 10 pCd and 12 pC1 neurons in female brains. Based on neurite projection patterns and anatomical location, these cells were postsynaptic to *DvPDF-Gal4*-expressing circadian neurons (Fig. 4B-C). Restricted *trans*-TANGO validation used sparse LexA drivers to target discrete *dsx*-positive neuronal subsets; these data indicated that pC1 and pCd but not pC2 are directly downstream of *DvPDF-Gal4*-expressing neurons (Fig. 4D). Although there were only 1-3 labeled cells per hemisphere, these drivers probably label only a specific fraction of their respective cell types (*83*).

Connectivity analysis of the circadian LNds reveals strong asymmetry (Fig. 5). Only the six Cry-positive LNds (LNd_a and LNd_b or E1 and E2) are extensively interconnected within the clock network (*10*, *14*). In contrast, the six cry-negative LNds (LNd_c or E3) are poorly connected to other circadian neurons but have extensive output connections to dsx-expressing downstream neurons (Fig. 5A-B). Specifically, LNd_c subtypes contribute disproportionately high synaptic and per-cell output compared to LNd_a/b (Fig. 5C-D). In males, LNd_c subtypes show robust and repeated connectivity to pCd-1, pC1b and pC1c neurons, with multiple LNds converging onto the same dsx-positive target neurons (Fig. 5A, 5C, 5E). While these contacts are less extensive in females, they remain the dominant circadian input to pCd-1 and pC1b neurons (Fig. 5B, 5D, 5E). These analyses therefore indicate that the six cry-negative LNds function less as internal clock integrators and more as specialized linkers of circadian neurons to sexually dimorphic outputs, with both synaptic strength and the number of engaged downstream neurons exhibiting sex-specific differences.

**Figure 5.**
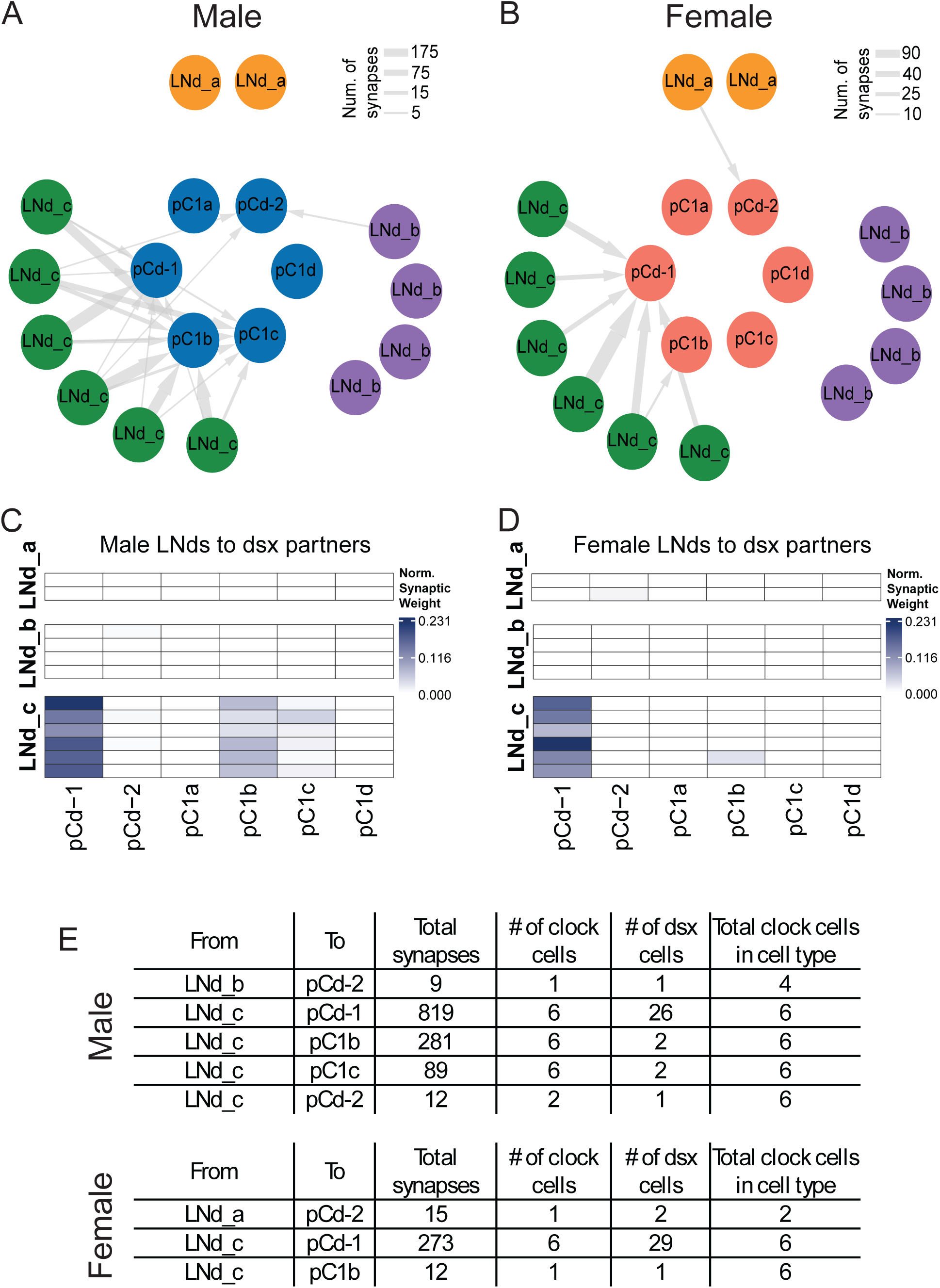
Sex-specific synaptic connectivity between LNd neurons and dsx-expressing downstream partners. **(A-B)** Network representation of synaptic connectivity from male **(A)** and female **(B)** LNd neurons to dsx-expressing downstream partner cell types. Nodes represent individual LNd neurons (Cry-positive: LNd_a and LNd_b, Cry-negative: LNd_c) and dsx partner cell types. Edges indicate synaptic connections; edge width is proportional to the total number of synapses between an individual LNd and a given partner cell type. Only connections with at least 5 synapses are shown. **(C-D)** Heatmap showing normalized synaptic output from male **(C)** and female **(D)** LNds to dsx-expressing cell types. Rows show LNd subtypes (Cry-positive: LNd_a and LNd_b, Cry-negative: LNd_c), and columns correspond to downstream partner groups. For each LNd neuron, synaptic weights were calculated by summing the total number of output synapses to all partners of a given cell and normalizing to the total number of output synapses from that neuron. Color indicates the total synaptic output of each neuron allocated to each partner group. **(E)** Summary tables of absolute synapse count underlying the network and heatmap representations. Columns indicate the total number of synapses for each LNd-partner pairing, the number of LNd neurons contributing to the connection, the number of dsx-expressing downstream neurons involved, and the total number of each LNd per subtype. Male (top) and female (bottom) datasets are shown separately.

To test whether this anatomical linkage represents a functional circuit, we performed optogenetic stimulation paired with calcium imaging (Fig. 6A) (*24*). Briefly, a red-light-activatable channelrhodopsin (UAS-CsChrimson-td-Tomato) was expressed under *DvPDF-Gal4* control, and *dsx*-LexA promoted the expression of a calcium indicator (LexAop-GCaMP6s). In both male and female brains, intermittent red-light stimulation of DvPDF-labeled neurons resulted in an increase in GCaMP signal from the cell body and projections of *dsx*-positive neurons (Fig. 6B-E), indicating that the circadian LNds have functional connections to *dsx*-positive pCd and pC1 neurons.

**Figure 6.**
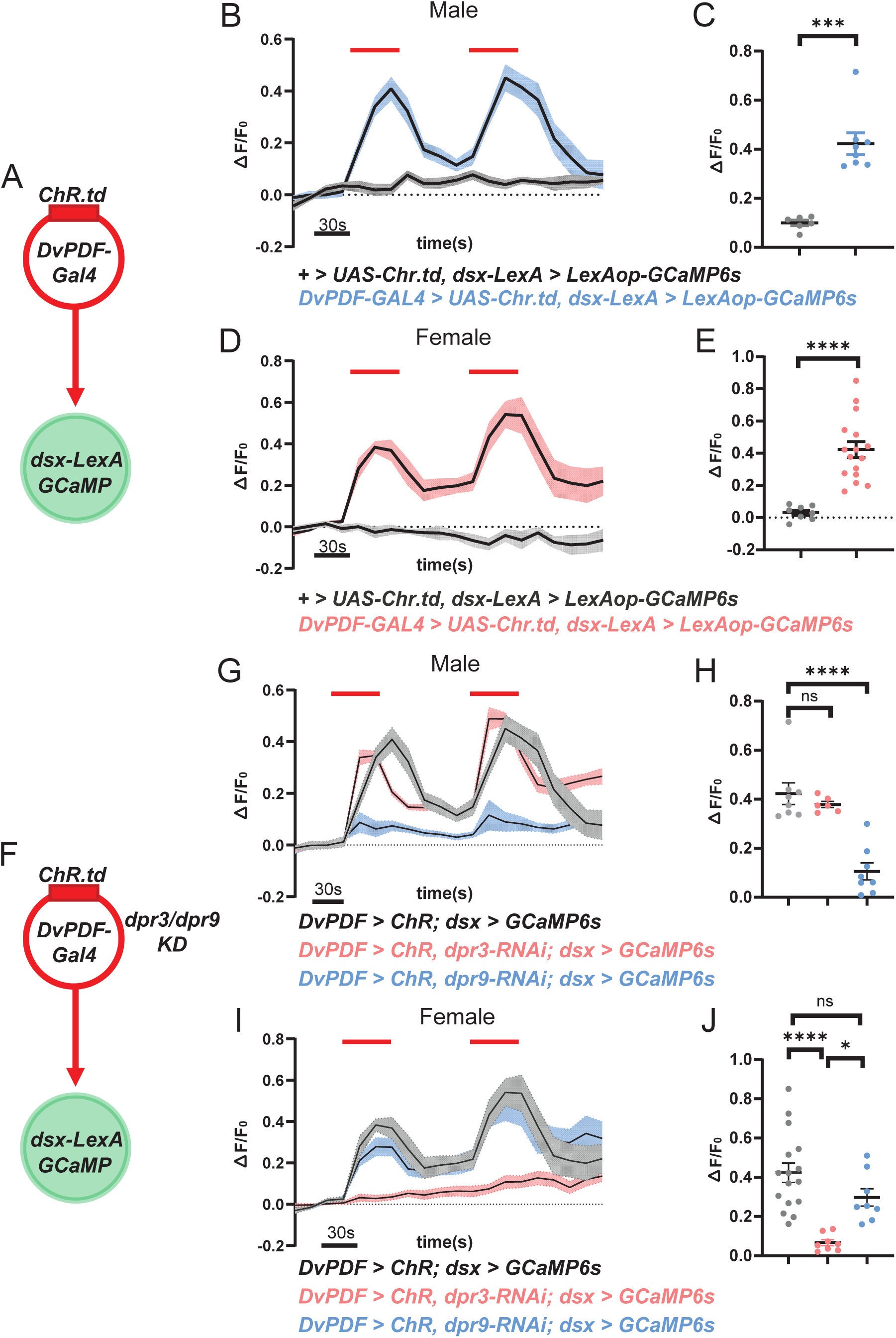
Sex-enriched CAMs in the LNds mediate synaptic connection to dsx-positive neurons. **(A, D)** Schematic representation of live imaging dsx-expressing neurons containing a Ca2+ indicator upon optogenetic activation of *DvPDF*-expressing neurons. **(B-C)** Male **(B)** and female **(C)** Ca2+ indicator traces of dsx-expressing neurons. X axis shows time; Y axis shows fluorescence intensity normalized to background signal. The red horizontal line represents the time and duration when red light was turned on for optogenetic activation. Quantification of these responses are shown on the right. Pink lines and dots show female samples, while blue dots show male samples. **(E-F)** Male **(E)** and female **(F)** Ca2+ indicator traces of dsx-expressing neurons upon a knockdown of sex-specific cell adhesion molecules in *DvPDF*-expressing neurons. Pink lines and dots represent knockdown of the female-enriched CAM, dpr3. Blue lines and dots represent knockdown of the male-enriched CAM, dpr9, in the LNds. Quantification of these responses are shown on the right. The red horizontal line represents the time and duration when light was turned on for optogenetic activation of *DvPDF*-expressing neurons.

Given that CAMs are known targets of sex-specific *fru* isoforms and drive sex-specific projection patterns of neurons (*41*, *84*, *85*), we asked whether *dpr9* and *dpr*3, the top sex-enriched CAMs in males and female LNd_c respectively, contribute to functional connectivity between these neurons and their downstream *dsx*-positive targets. To this end, we performed genetic knockdowns of either *dpr3* or *dpr9* under *DvPDF-Gal4* control while recording GCaMP6s signals from the cell body and projections of *dsx*-LexA labeled cells (Fig. 6F).

Knockdown of the male-enriched CAM, *dpr9*, in male LNds resulted in a significant reduction of GCaMP signal in downstream *dsx* neurons. In contrast, knocking down female-enriched *dpr3* in male cells had no effect, i.e. it had comparable GCaMP signal to the control condition (Fig. 6G, H). Similarly, knocking down the female-enriched CAM, *dpr3*, in female LNds resulted in a significant reduction in GCaMP signal, whereas the *dpr9* knockdown in female LNds resulted in a modest, non-significant decrease in Ca^2+^ influx (Fig. 6I, J). These results indicate that these sex-enriched CAMs in the LNds are essential for establishing or maintaining functional connectivity to downstream sexually differentiated neurons.

## Discussion

The recent transcriptomic profiling of *Drosophila* clock neurons has unveiled dramatic transcriptional heterogeneity within the circadian circuit, far exceeding what anatomical classifications had suggested (*25*). However, these comprehensive transcriptomic maps had ignored the extent to which sex might contribute to transcriptomic differences. This possibility is also interesting because circadian neurons and clock-controlled genes are involved in sexually dimorphic physiologies and behavior. For these reasons, we characterized the sexual dimorphism of *Drosophila* clock neurons. The results include at least one DN1p and two DN3 clusters, the latter being arguably the most heterogeneous and enigmatic group of clock neurons (*10*, *25*, *26*, *53*). There is also a sexually dimorphic group of LNds: the E3 LNds. These neurons not only have highly sexually dimorphic transcriptomes but also connect to some of the central regulators of sexually dimorphic behaviors.

In our first transcriptomic atlas of clock neurons (*25*, *26*), we identified two LNd clusters that we interpreted as distinct subtypes; however, our present analysis reveals that this subdivision reflects a single LNd subtype split by sex. In contrast, the other sexually dimorphic clusters identified in this study did not segregate in the original dataset. Several technical and biological factors may explain this difference. First, the present work isolated clock neurons using an H2A-GFP fusion protein, which is stronger and more stable than the cytoplasmic eGFP reporter used previously. The current work may therefore have labeled a broader population of circadian neurons, increasing our ability to detect subtle transcriptomic differences in the adult brain. Second, the earlier dataset may have recovered relatively few neurons of each sex per time point, potentially limiting statistical power to resolve sex-specific transcriptional signatures. As a result, clusters that are now clearly dimorphic may have previously appeared as single, mixed-sex populations.

The hierarchical organization of the LNds positions them as core activity-promoting neurons; the E1 and E2 LNds are important in different aspects of locomotor activity behavior (*14–16*). The E3 cells in contrast have no known role in these behaviors. Moreover, circadian connectivity analyses have identified only sparse interactions between the E3 LNds and the rest of the circadian circuitry, suggesting that these neurons may predominantly connect to neurons that are upstream or downstream of the clock neurons. For example, the E3 LNds may connect to inputs that help adapt the clock to temperature (*14*, *86*). Additionally, studies have implicated the LNds in sexually dimorphic behaviors such as rival-induced mating duration, mating drive, and oviposition. However, the roles of specific LNds like the E3 LNds in these behaviors has not been previously shown despite evidence that the clock is involved and that the expression of *per* and *tim* in circadian neurons is required for rival-induced long mating duration (LMD) (*87*).

A key question arises from our findings: since the E3 LNds are relatively isolated from communication to many other clock neurons but maintain robust connections with pC1 and pCd neurons in the adult, what signals are they relaying? Although both male and female E3 LNds form substantial synaptic outputs onto pC1 and pCd-1 neurons, synaptic connections in the opposite direction (dsx-to-LNds) are far fewer, consistent with a model in which this circuitry predominantly conveys timing information from LNds to sexually dimorphic neurons. Neuronal connectivity molecules (e.g., CAMs) are known to regulate wiring specificity, and sexually dimorphic expression of distinct *dpr* genes in male and female LNd_c contribute to differences in their downstream connectivity to dsx-expressing neurons, including the male-specific connections to pC1c (Fig. 5). Our results further suggest that multiple Dprs can converge onto the same target neuron. Although no female-specific dsx-positive target neurons were identified, *dpr3*-RNAi in female LNds abolishes functional connectivity with dsx-positive neurons (Fig. 6I-J). Perhaps pCd-1 neurons, the primary sex-shared targets of LNd_c (Fig. 5), contain a heterogeneous population, with distinct subtypes targeted by male and female LNd neurons.

We did not observe locomotor activity defects upon a *dpr9* or *dpr3* knockout (KO) in the LNds. Surprisingly, CRISPR-mediated knockout (KO) of several candidate LNd dimorphic genes, including those described above, also had no significant effect on mating activity, which included mating latency and mating duration (data not shown). These negative results may have been due to redundant or even compensatory molecules that can functionally substitute for these *dprs*. Alternatively, the impact of these CAMs may be too subtle to detect in standard behavioral assays, these molecules may regulate other sexually dimorphic behaviors not addressed in our current assays, or the circuitry may be sufficiently resilient or buffered to sustain basic mating behaviors under laboratory conditions despite the fact that the two CAMs modulate the functional strength of the LNd-to-pC1/pCd connections (Fig. 6).

In *Drosophila*, the earliest detection of *fru* transcripts coincides with the first appearance of what are thought to be the circadian LNd and DN3 neuronal clusters. The sexual dimorphism of these neurons likely originates from the late stages of pupal development (*53*, *88–90*). It is possible that, as these *fru*-expressing neurons develop, sex-specific isoforms of FRU regulate the expression of sexually dimorphic connectivity molecules, which influence gene expression and developmental wiring in a sex-dependent manner. This could explain why the misregulation of genes involved in sexual differentiation results in altered locomotor activity behaviors in both sexes.

Neuronal cell number is also thought to be regulated by *dsx* and *fru* expression. Indeed, we find here that some *fru*-expressing circadian neuron populations vary in the number of male and female cells detected. The two transcriptionally related DN3 clusters share many defining features that distinguish them from other clock neurons but do not exhibit the same trend in neuron numbers. Although a previous study on *fru* regulation determined that distinct *fru* isoforms affect DN3 cell numbers (*51*), it is not known how FRU influences neuronal cell numbers. Recent findings also suggest that the DN3 population contains sleep-promoting neurons (*25*). While that study did not identify specific DN3 subclusters that promote sleep, it suggests that it will be interesting to identify the role of the sexually dimorphic DN3 subclusters shown here in circadian-influenced behaviors.

GPCRs and neuropeptides are major contributors to neuronal communication. Unlike traditional synaptic communication, GPCR and neuropeptide connections probably do not manifest in connectivity analyses. Nonetheless, these molecules relay signals to neuronal partners to enable the proper execution of physiological or behavioral outputs. Our lab has previously shown that GPCRs and neuropeptides alone can define adult circadian neuron populations (*27*), and the present study further shows that these connectivity molecules are also sex-enriched within a cell type. These findings suggest that sexually dimorphic GPCRs and neuropeptides contribute to aspects of sexually dimorphic circadian circuits not only during development but also in the fully developed adult nervous system.

Further, social enrichment influences mating duration, and GPCR/neuropeptide signaling within circadian lateral neurons is required for this context-dependent mating behavior (*77*). Future studies will likely determine additional roles for these sexually dimorphic GPCRs and neuropeptides in social context-dependent behaviors.

In summary, our work expands upon previous studies by adding the dimension of sex to the transcriptomic profile of *Drosophila* circadian neurons. Beyond identifying sexually dimorphic circadian clusters and genes, our study offers functional insights into their underlying connectivity. Understanding more about the regulation of sexually dimorphic gene expression in clock neurons will clarify how gene regulation interacts with environmental factors to influence behavioral outputs of the clock network.

## Material and Methods

### Reanalysis of published single-cell RNA-seq data set

Single-cell transcriptomic sequencing data were processed using cellranger version 7 (*91*). Files were processed individually to obtain FASTQ files and mapped to a genome containing lncRNAs (dm6). Mapped reads to lncRNAs in each cell were plotted in Fig. S3A. Data integration was performed using Seurat V4 (*92*); a merged RDS object containing the six single-cell experiments (one for each timepoint) was integrated using time as a common variable; this was done via FindIntegrationAnchors and IntegrateData (*92*). This ensured that the clusters generated were not influenced by cycling gene expression. *de novo* cluster identification for the remapped dataset was performed by comparing the average gene expression profile of each newly identified cluster to those previously established (*26*, *25*). Additional validation of cluster identity was done by Pearson’s gene expression correlation between the remapped dataset and the original dataset (Fig. S3) (*25*).

### Fly strains and husbandry for single-cell RNA-seq experiments

Flies were reared on standard cornmeal and agar medium with yeast under 12:12 LD conditions at 25°C. Females used for single-cell experiments were collected as virgins and paired with males of the same genotype 3 days post-eclosion. Females were co-reared with males for 3 days. A list of flies used in this study can be found in Table S1.

The new single-cell dataset was generated using a genetic multiplexing technique that enables pooled processing of all circadian time points. This approach has previously been used to generate developmental time-series datasets with high temporal resolution (*93*, *94*). Virgin female flies expressing the Clk856-Gal4 driver were crossed to male flies expressing a nuclear GFP reporter (UAS-His2A-GFP) and carrying one of twelve unique 3rd chromosomes from wild-type strains of the Drosophila Reference Genetic Panel (DGRP) (*95*). Each DGRP chromosome carries a unique set of known single-nucleotide polymorphisms that can be used to separate cells from individual crosses (*96*). F1 flies were collected 1-4 hours after eclosion; sexes were raised separately for 3 days. On day 4, male and female flies with the same DGRP genotype were combined and entrained for 4 days in 12:12 LD. Flies were collected at ZT02, ZT06, ZT10, ZT14, ZT18, and ZT22. Two unique DGRP genotypes were used as biological replicates at each time point, and equal numbers of male and female flies were collected for each genotype. Flies from all time points and DGRP genotypes were combined and processed as a single sample from brain dissections to single-cell dissociation and processing. Two independent samples were generated using this approach.

For the sex-informed data set, virgin female flies expressing Clk856-Gal4 driver and UAS-eGFP were crossed to males expressing a split Gal4 that labels DN3 (R11B03-AD; VT002670-DBD) (*25*). F1 progenies were collected 1-4 hours after eclosion. The sexes were raised separately and entrained in 12:12 LD for 3 days. Following entrainment, male and female flies were combined for 2 days and later separated for 3 days in the same LD conditions. Male and female flies were collected separately at ZT06 and ZT18 and put on ice before dissection.

Approximately 200 brains were dissected in ice-cold Schneider’s *Drosophila* medium (SM, Gibco #21720001) and then briefly washed with SM and incubated at room temperature with 0.75ug/ul Collagenase (Sigma-Aldrich #C1639) and 0.04 μg/ul Dispase (Sigma-Aldrich #D4818) for 30 minutes. Brains were washed twice with fresh SM, mechanically triturated to obtain a single cell suspension, and passed through a 40um filter. The volume was adjusted to 3mL for FACS sorting. Nucblue (Invitrogen #R37605) was used to stain for living cells, and GFP was used to isolate the GFP-positive clock neurons via a fluorescence-activated cell sorter (FACS, BDFACS Melody). FACS gates were set using neurons that did not express GFP. 35,000-40,000 GFP-positive cells per replicate were sorted into 500uL of 1XPBS + 0.5% BSA.

### Generation of single-cell libraries and raw data processing

Isolated cells were spun down at 1,000rcf for 10 minutes at 4°C using a swing bucket adapter and loaded in the GEM chip from the Chromium Single Cell 3′ Kit (version 3 of 10X Genomics). RNA-seq libraries were prepared using the manufacturer’s recommended protocol (CG000315 Rev. B; 10X Genomics). Paired-end sequencing was performed using the Illumina NovaSeq 6000 at 150 cycles. The 10X Cell Ranger package was used to align the sequencing data to the *Drosophila* genome (dm6), including long non-coding RNAs. Reads aligned to annotated exons were quantified using unique molecular identifiers (UMIs) to quantify RNA molecules accurately.

For the new single-cell data set with DGRP chromosomes, single-cell transcriptomes from different timepoints and replicates were demultiplexed based on the unique genotypes of the DGRP chromosomes (*95*). Demultiplexing was performed using demuxlet (*96*) as described in Dombrovski et al. (*94*)

### Data integration and clustering analysis

To retain only high-quality single-cell transcriptomic data, we filtered out cells based on several criteria: 1) the number of detected genes had to be between 500 and 3000, 2) the number of detected UMIs had to be between 1000 and 25000, and 3) the cells with a percentage of mitochondrial RNA greater than 7%.

Clustering analysis of the new sex-inferred data set was done using the Seurat package (*92*). High-quality cells from two independent replicates were separated by DGRP line and time using SplitObject, data was normalized, and 2000 variable features were identified using NormalizeData and FindVariableFeatures, respectively. These variable features were used to integrate the datasets using FindIntegrationAnchors and IntegrateData. Clusters were generated using time as a constant to avoid clustering based on time-of-day gene expression. After applying our standard Seurat clustering pipeline (*92*), we removed only clusters with low *tim* or *Clk* gene expression (for *tim* less than 5 TP10K, for *Clk* less than 1 TP10K), confirmation was done by observing cycling expression of these genes. Cluster identification and naming was performed by comparing known marker gene expression from the individual clusters to those established in previous studies (*26*, *25*). Processing and clustering of the sex-informed data set were performed similarly to the procedures described above. Briefly, the data sets were separated into time points, 2000 variable features were identified, and the common features were used to generate the clusters. Because there is no time course in this experiment, the cycling expression of core clock genes was not used to subset out circadian clusters. Instead, marker gene expression was compared to previously defined clusters (*25*) and confirmed by a correlation matrix between the sex-informed and sex-inferred data sets.

### Differential gene expression and Gene Ontology analysis

To identify sex-enriched features within the circadian cell types, we first divided the cells into cell type and sex. Using this categorization, we applied FindMarkers from the Seurat analysis package to compare the transcriptomic profile between the sexes for each cell type (*92*). Sex-enriched transcripts were defined using an adjusted p-value of < 0.05 and a fold change of 1.7 or greater.

Gene Ontology enrichment analysis was performed using PANGEA (*97*) with genes expressed in all clock neurons as a background gene set. Gene sets analyzed included the differentially expressed genes in males or females from the top 4 dimorphic clusters. Log2 normalized counts were used to generate heatmaps illustrating the expression of cell surface molecules, GPCRs, and neuropeptides. Heatmaps were created using the ComplexHeatmap package in RStudio (*98*).

### Immunohistochemistry

Flies were fixed by rotating them in 1X PBS with 4% paraformaldehyde (PFA) and 0.5% Triton X-100 for 2.5 hours at room temperature. Following fixation, flies were washed 3X with 1X PBS and 0.5% Triton X-100 (PBST) for 10 minutes each wash at room temperature. Brain dissections were done in 1X PBS solution and blocked with 10X normal goat serum (NGS; Jackson Labs 005-000-121) for 1 hour at room temperature or overnight at 4°C. This was followed by an overnight incubation with primary antibodies against GFP (chicken anti-GFP; Abcam ab13970) and RFP (rat anti-RFP; ProteinTek #5f8) at 4°C. Brains were washed 3X for 15 minutes at room temperature with 0.5% PBST prior to being incubated with secondary antibodies (Alexa Fluor 488 conjugated anti-chicken, Alexa Fluor 546-conjugated anti-rat diluted at 1:500 in 10X NGS + 0.5% PBST for 2 hours at room temperature. Brains were washed with 0.5% PBST 3X for 15 minutes. Brains were mounted in VectaShield mounting media (Vector Laboratories H-1900-10) and imaged using a Leica Stellaris 8 confocal microscope with a 20X dry or 40X oil objective. Images were processed using FIJI (*99*).

### Connectome Analysis

Synaptic connectivity data for LNd neurons were obtained from the female (FAFB v783) and male (MCNS v0.9) connectome (*28*, *29*, *100*). For each individual LNd neuron, synaptic connectivity was quantified by summing the total number of synapses between that neuron and all downstream partners belonging to the same cell type. The connectivity network was visualized using Cytoscape (*101*). Because the male and female connectomes differ in synaptic annotation density, with higher overall synaptic resolution in the male dataset, direct quantitative comparison of absolute synapse numbers between sexes should be interpreted with caution (*28*, *29*, *100*). To account for differences in total synaptic output across individual LNds, synaptic weights were normalized to the total number of output synapses for each neuron. These normalized synaptic weights were visualized as heatmaps, with rows corresponding to individual LNd neurons and columns corresponding to downstream dsx-expressing partner cell types. Heatmaps represent the fraction of each neuron’s total synaptic output allocated to each partner group.

### Two-photon Functional Calcium Imaging

For *ex vivo*, adult flies were fed with ATR for 3-5 days and dissected during the ZT4-8 period. The fly brains were dissected in hemolymph-like saline (AHL) containing 108mM NaCl, 5mM KCl, 2mM MgCl_2_, 2mM CaCl_2_, 4mM NaHCO_3_, 1mM NaH_2_PO_4_, 5mM HEPES, 10mM Sucrose, 5mM Trehalose, pH 7.2. The dissected brain was mounted on a special dish containing 1 mL of AHL to prevent movement during imaging. To activate the specific neurons expressing the CsChrimson, we used 630nm red LED illumination (∼0.1mW*mm-2) through an optical fiber to illuminate the brain in the dish. For two-photon Ca2+ imaging, we used an Olympus FV1200MPE multiphoton laser scanning microscope equipped with a 25x 0.8NA objective (GCaMP7s excitation @920 nm). Recordings lasted ∼120 seconds (two Opto-Stim at 20-35s and 60-75s). Imaging parameters varied slightly between experiments. Images were typically acquired at a resolution of 512×512 pixels and a frame rate of approximately 0.9 Hz. ROIs were analyzed with Fiji. Fluorescence change was calculated using the formula:△F/F_0_ = (F_t_-F_0_)/F_0_×100% and △FMax/F_0_ = (F_t_Max-F_0_)/F_0_×100%, where F_t_ is the fluorescence at time n, and F_0_ is the fluorescence at time 0.

### Generating transgenic flies

#### Ca^2+^ indicator fly line for functional connectivity

The QUAS-jGCaMP7b plasmid was constructed through molecular cloning procedures at Fungene Biotech (http://www.fungene.tech). The coding sequence of jGCaMP7b was PCR-amplified from the 20XUAS-IVS-jGCaMP7b flies (Bloomington Stock Center #79029) using high-fidelity DNA polymerase. The amplified product was subsequently cloned into the pQUAST expression vector (Addgene Plasmid #24349). Finally, the recombinant plasmid was integrated into the VK00005 genomic landing site using site-specific integration techniques to generate stable transgenic lines.

