## Supplemental Figures for "Transcriptomic and functional characterization indicate sexual dimorphism of discrete circadian neuron subtypes"

**Figure S1. Validation of sex-informed *de novo* cluster identification and sex-assignment cutoffs**

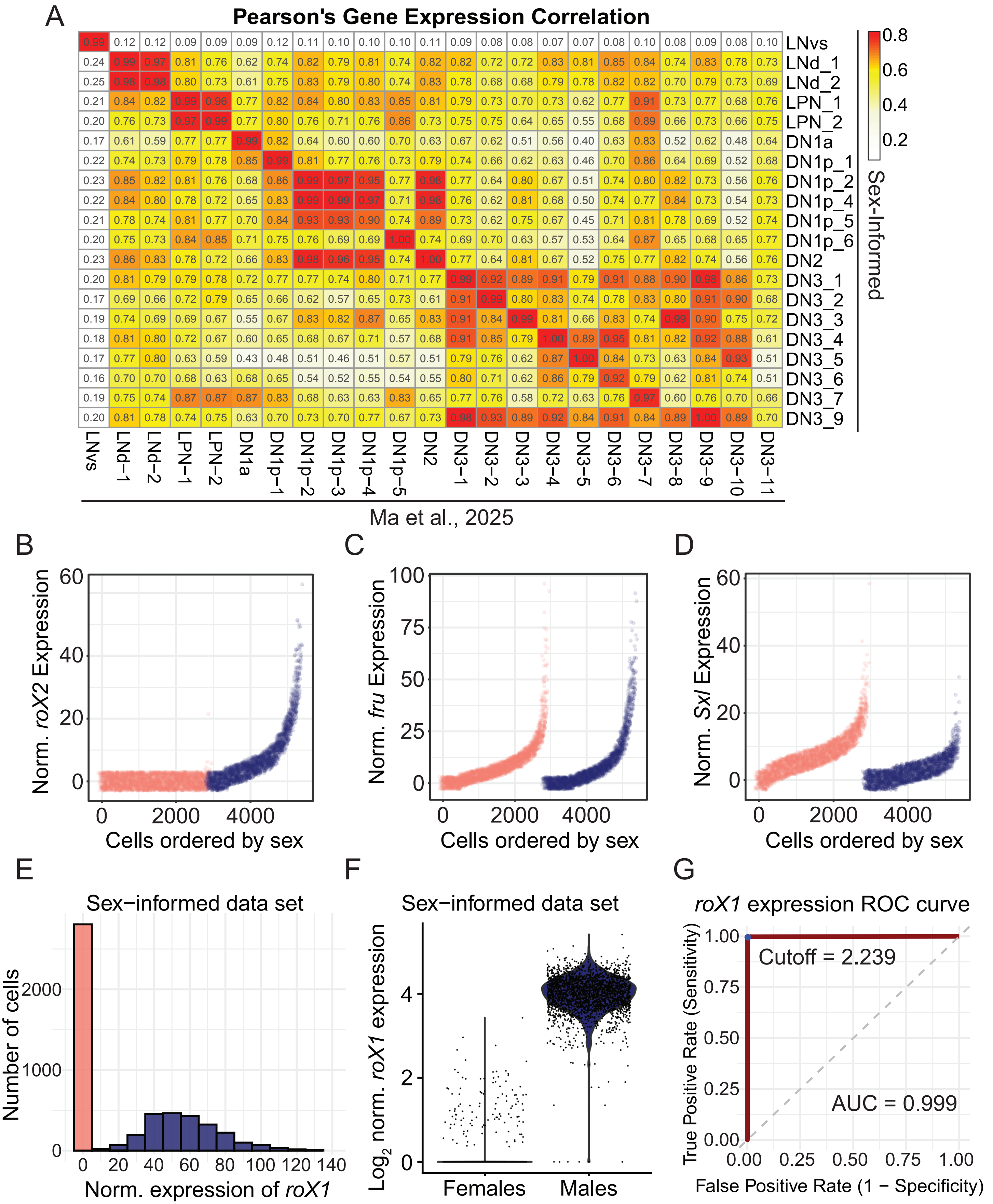

**Figure S2. Validation of sex-assignment in sex-inferred dataset based on sex-informed cutoff**

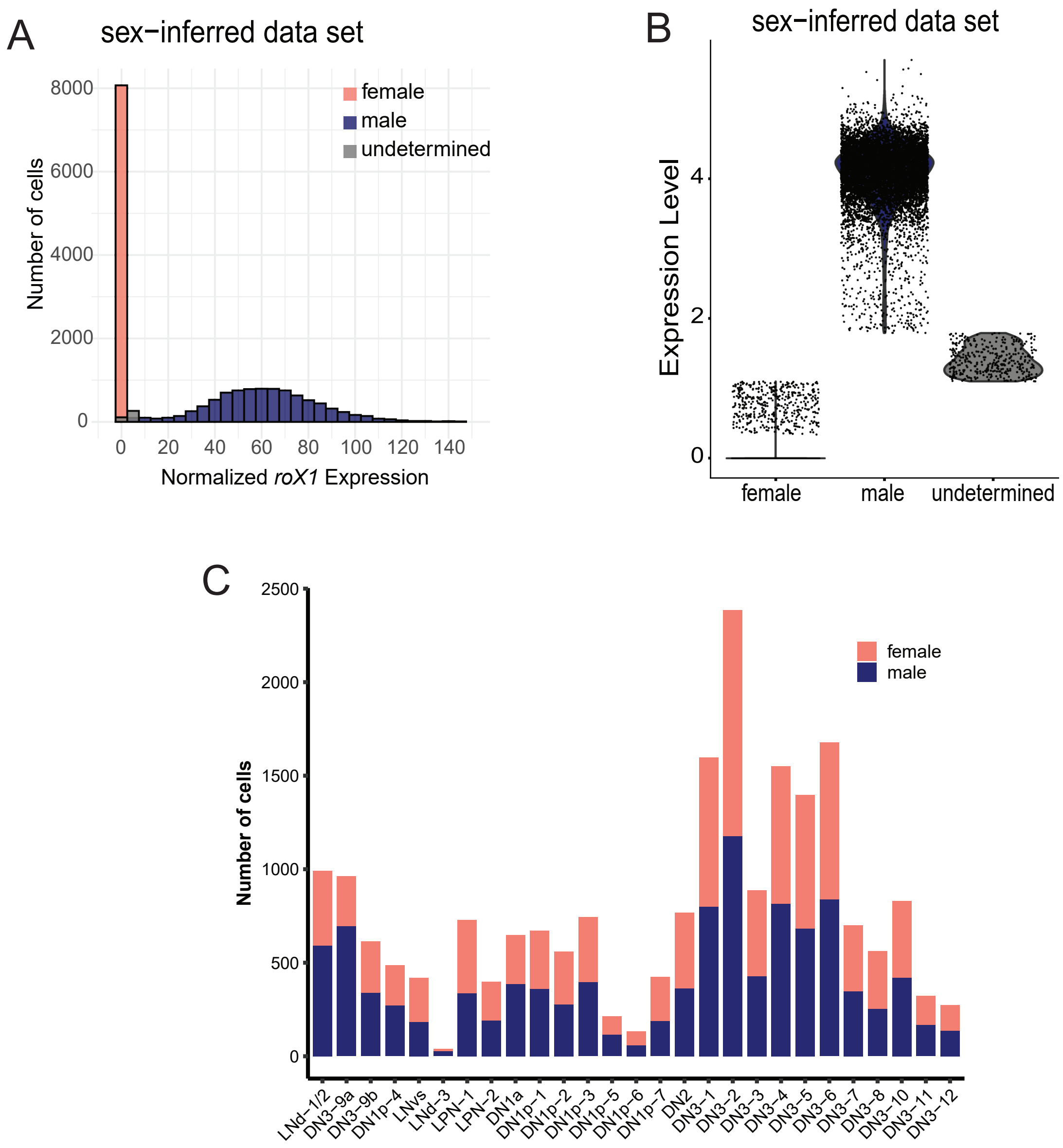

**Figure S3. Remapping and reclustering previously published circadian single-cell datasets**

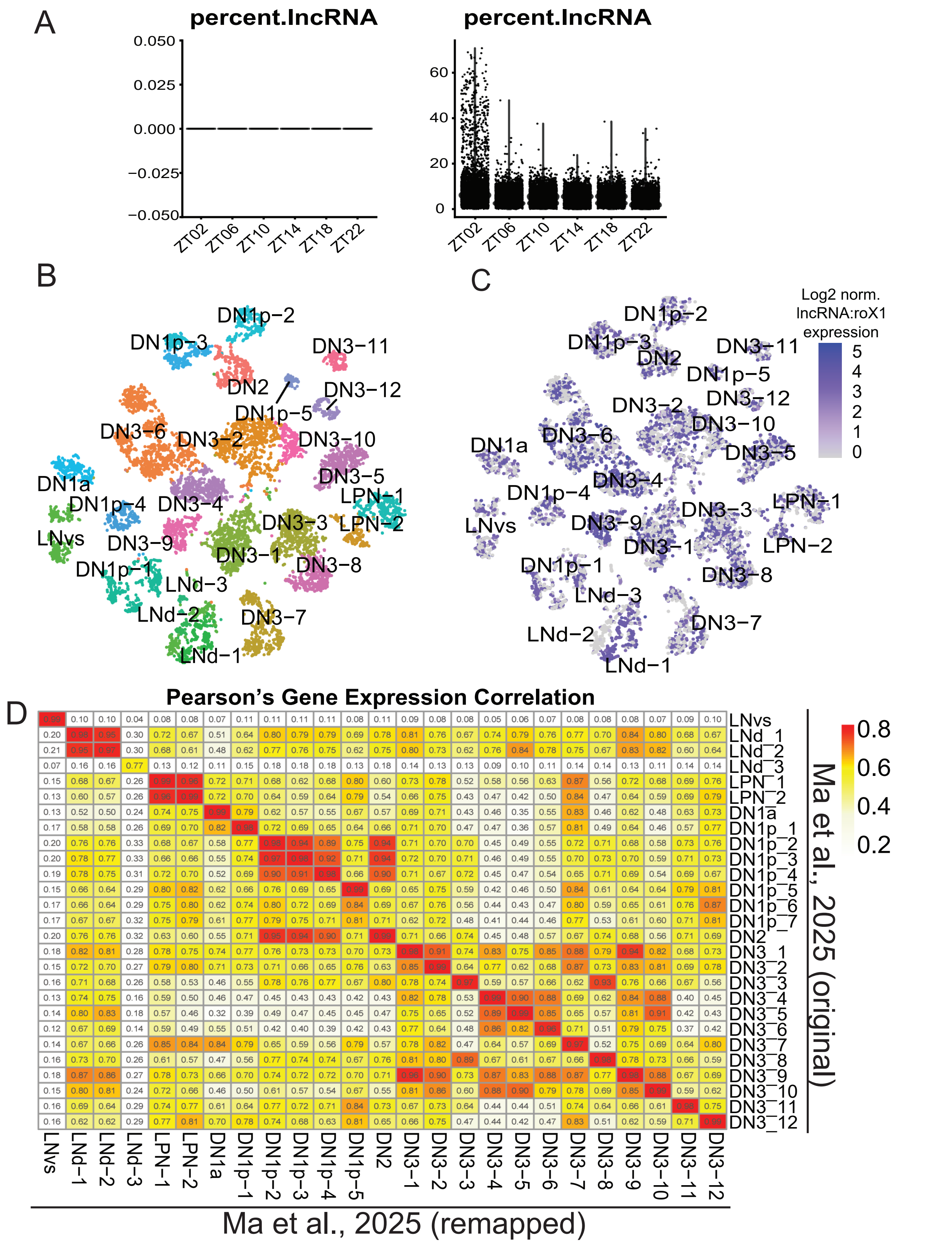

**Figure S4. Marker gene expression in the sexually dimorphic clusters**

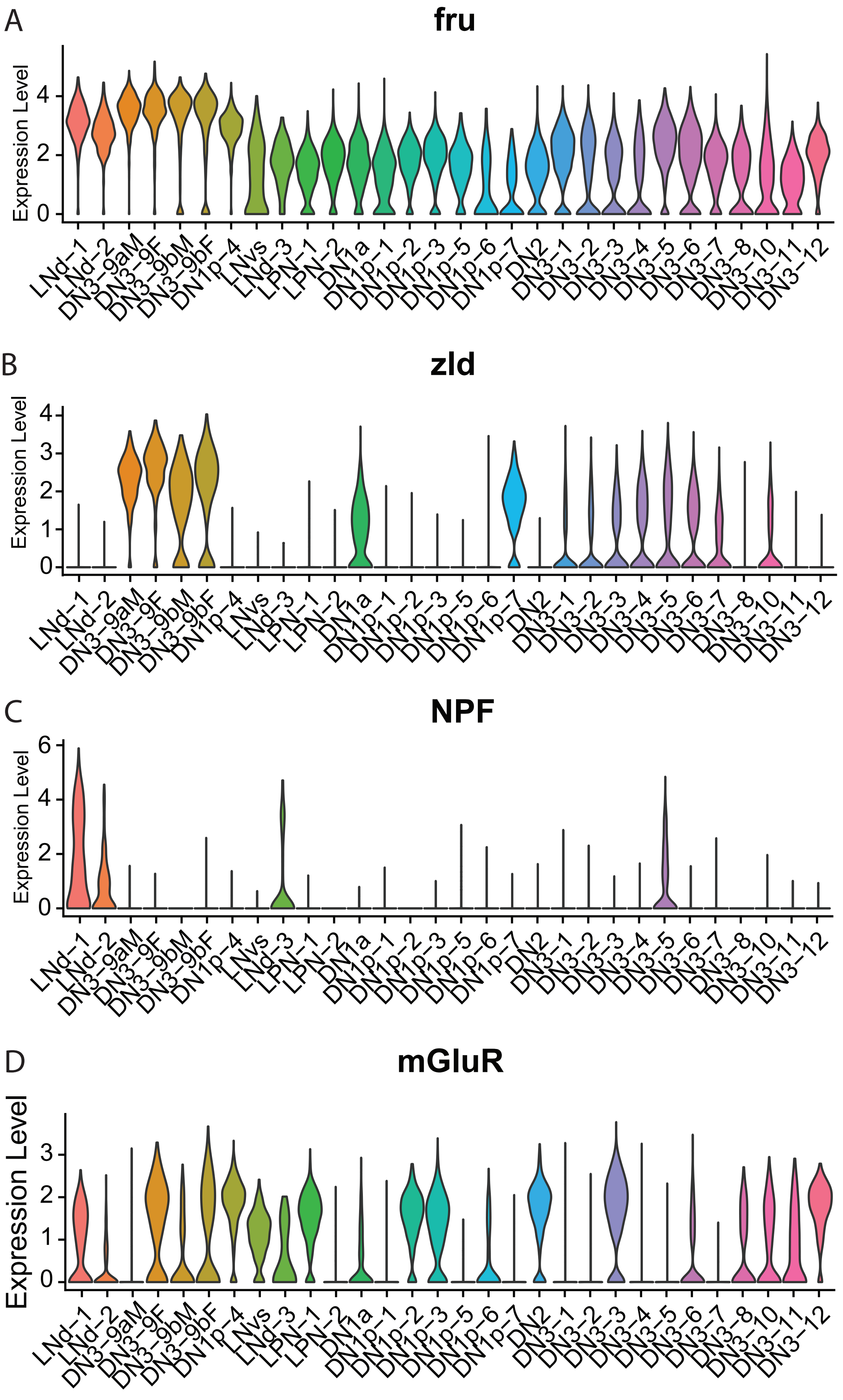

**Figure S5. Sex-specific neural connectivity molecules within the DN3 dimorphic cell types**

DN3-9a

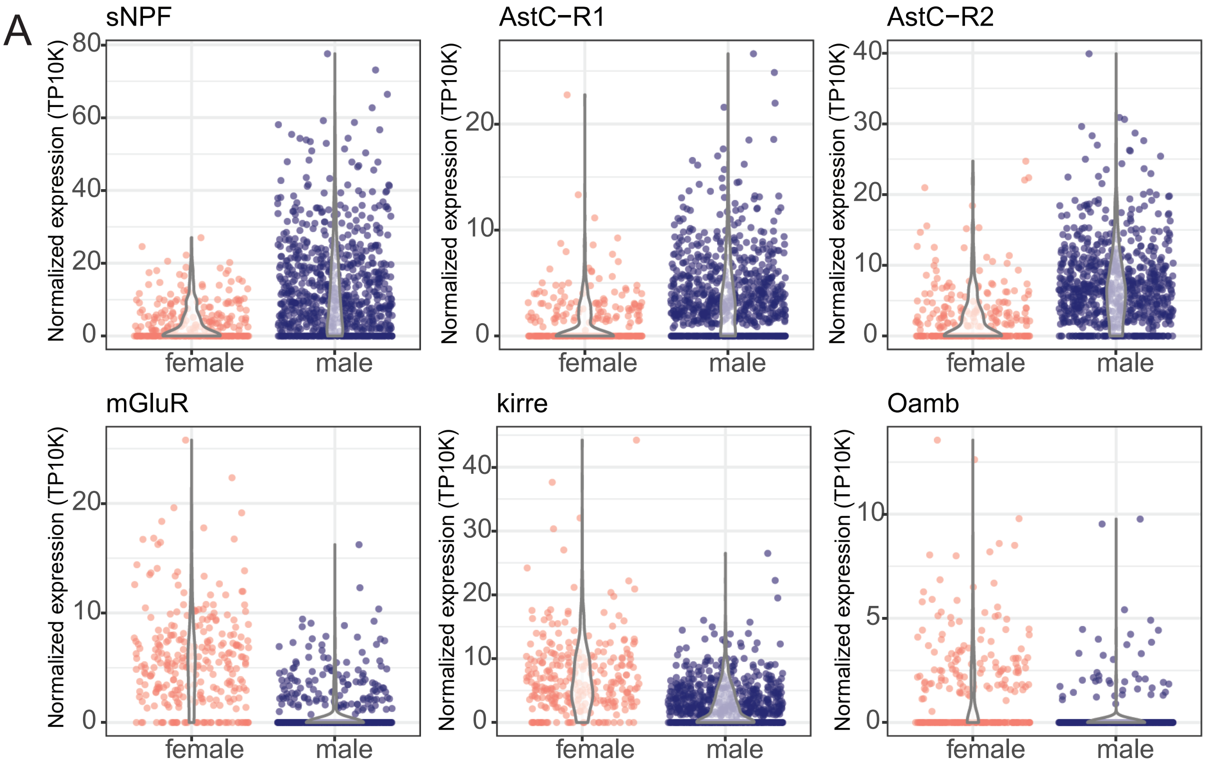


DN3-9b

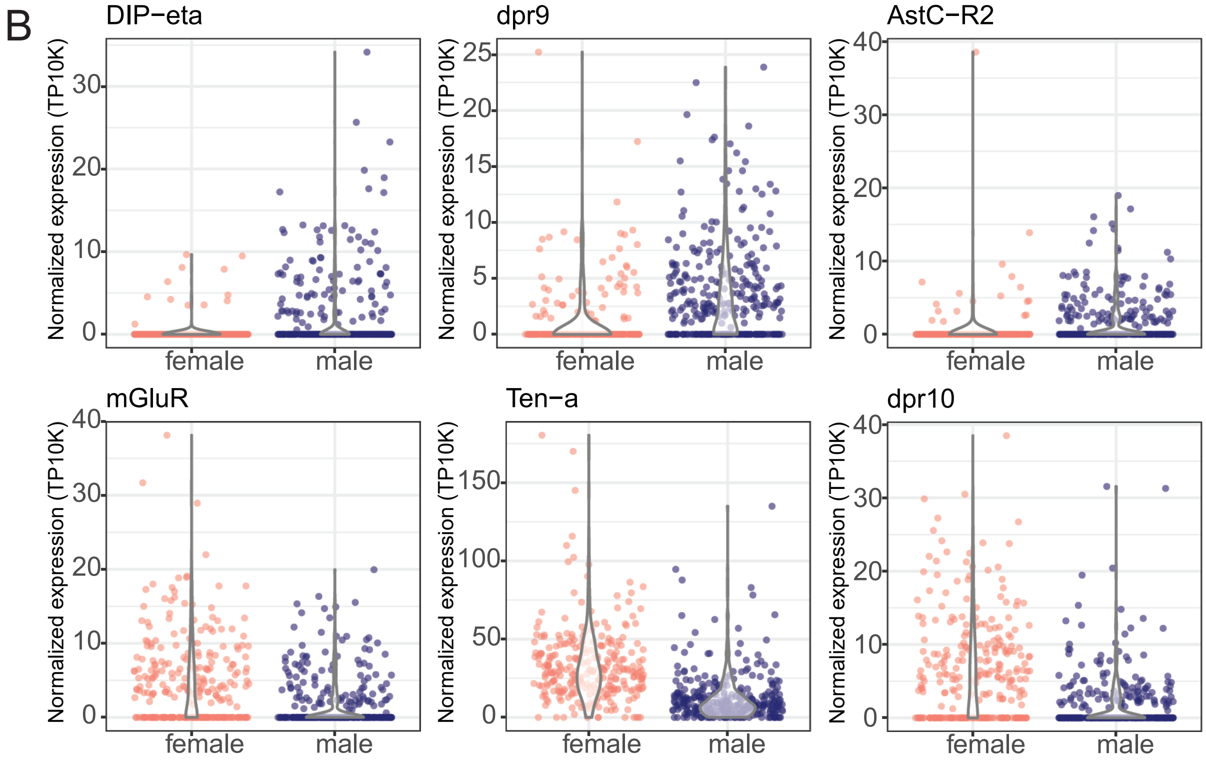

Figure S1. Validation of sex-informed *de novo* cluster identification and sex-assignment cutoffs.

**(A)** Pearson's gene expression correlation heatmap between the newly generated sex-informed dataset and previously established circadian clusters from Ma et al., 2025. Numbers close to 1 show high correlation between the newly identified clusters and established cluster identity. **(B)** Normalized expression of *lncRNA:roX2* in sex-informed data set. Pink dots represent female-derived cells, blue dots show male-derived cells. **(C)** Normalized expression of *fru* in sex-informed data set. Pink dots represent female-derived cells, blue dots show male-derived cells. **(D)** Normalized expression of *Sxl* in sex-informed data set. Pink dots represent female derived cells; blue dots show male-derived cells. **(E)** Histogram plot of male and female derived cells against normalized *lncRNA:roX1* gene expression. Pink bars represent number of female cells; blue bars show the distribution of male cells with different *lncRNA:roX1* expression levels. **(F)** Log2 normalized *lncRNA:roX1* gene expression of male and female derived cells in sex-informed data set. Each dot represents a cell, pink violin shows female cells, blue violin show male cells. **(G)** AUROC plot used to determine optimal *lncRNA:roX1* expression cutoff based on sex-informed data set. X axis represents the false positive fraction; Y axis shows true positive fraction. Blue dot shows the optimal expression cutoff for calling a cell "male" (> 2.3) or "female" (< 2.3).

Figure S2. Validation of sex-assignment in sex-inferred dataset based on sex-informed cutoff.

**(A)** Histogram plot of male and female derived cells against normalized *lncRNA:roX1* gene expression. Pink bars represent number of inferred female cells; blue bars show the distribution of inferred male cells with different *lncRNA:roX1* expression levels. Grey bards show cells with undetermined sex based on stringent lncRNA:roX1 expression cutoffs. **(B)** Violin plot (normalized Log2 scale) of *lncRNA:roX1* gene expression of male and female derived cells in the sex-informed data set. Each dot represents a cell, pink violin shows female cells, blue violin show male cells. Grey violin show cells with undetermined sex based on stringent *lncRNA:roX1* expression cutoffs. **(C)** Number of cells per cluster separated by inferred sex. Pink bars show inferred female cells; blue bars show inferred male cells. Numbers represent the ratio of male to female neurons in the individual circadian clusters.

Figure S3. Remapping and reclustering previously published circadian single-cell datasets.

**(A)** Percentage of lncRNAs expressed in the single-cell data set from Ma et al., 2025. The original dataset did not include lncRNAs (left violin plot). Remapped dataset to a genome containing lncRNA shows a higher percentage of cells expressing lncRNAs (right violin plots). **(B)** Dimensional reduction plot showing cluster identities of remapped dataset. **(C)** Dimensional reduction plot showing log2 normalized *lncRNA:roX1* expression distribution in remapped dataset. Green dashed circles show *roX1* biased clusters. **(D)** Pearson's gene expression correlation of published single-cell dataset (*23*) and remapped dataset. Numbers closer to 1 reflect high cluster correlation indicating correct cluster identity assignment.

Figure S4. Marker gene expression in the sexually dimorphic clusters.

Log2 normalized gene expression of **(A)** *fru*, **(B)** *zld*, **(C)** *NPF*, and **(D)** *mGluR* in the circadian clusters. Dimorphic circadian clusters are shown on the left. Male-biased clusters are shown in blue, female-biased clusters are shown in pink. Clusters that show male and female co-clustering are shown in purple. *fru* is a defining feature of all clusters with dimorphic gene expression profile. *zld* is a defining feature of the dimorphic DN3 clusters (DN3-9a, DN3-9b). *NPF* and *mGluR* are defining features of male LNds. Bracket indicates sexually dimorphic circadian cell types.

Figure S5. Sex-specific neural connectivity molecules within the DN3 dimorphic cell types.

**(A)** Violin plot showing (TP10K normalized scale) expression of neural connectivity molecules upregulated in either male (top) or female (bottom) DN3-9a cells. CAMs (*kirre*), GPCRs (*AstC-R1, AstC-R2*, *mGluR* and *Oamb*) and neuropeptide (*sNPF*). **(B)** Violin plot showing (TP10K normalized scale) expression of neural connectivity molecules upregulated in either male (top) or female (bottom) DN3-9b cells. CAMs (*DIP-eta*, *dpr9*, *Ten-a*, *dpr10*), and GPCRs (*AstC-R2* and *mGluR*).
